## Supplementary material for "High frequency spike inference with particle Gibbs sampling": TeX source files: biblio.bib

@article{diana2019bayesian,
  title={Bayesian inference of neuronal assemblies},
  author={Diana, Giovanni and Sainsbury, Thomas TJ and Meyer, Martin P},
  journal={PLoS computational biology},
  volume={15},
  number={10},
  pages={e1007481},
  year={2019},
  publisher={Public Library of Science San Francisco, CA USA}
}

@article{chen2013ultrasensitive,
  title={Ultrasensitive fluorescent proteins for imaging neuronal activity},
  author={Chen, Tsai-Wen and Wardill, Trevor J and Sun, Yi and Pulver, Stefan R and Renninger, Sabine L and Baohan, Amy and Schreiter, Eric R and Kerr, Rex A and Orger, Michael B and Jayaraman, Vivek and others},
  journal={Nature},
  volume={499},
  number={7458},
  pages={295--300},
  year={2013},
  publisher={Nature Publishing Group}
}

@book{Chopin2020book,
    title={An Introduction to Sequential Monte Carlo},
    publisher={Springer-Verlag},
    year={2020},
    author={Chopin, N and Papaspiliopoulos, O}
}

@article{tubiana2020blind,
  title={Blind deconvolution for spike inference from fluorescence recordings},
  author={Tubiana, J{\'e}r{\^o}me and Wolf, S{\'e}bastien and Panier, Thomas and Debregeas, Georges},
  journal={Journal of Neuroscience Methods},
  volume={342},
  pages={108763},
  year={2020},
  publisher={Elsevier}
}

@article{hu2018detecting,
  title={Detecting T cell activation using a varying dimension Bayesian model},
  author={Hu, Zicheng and Lancaster, Jessica N and Ehrlich, Lauren IR and M{\"u}ller, Peter},
  journal={Journal of applied statistics},
  volume={45},
  number={4},
  pages={697--713},
  year={2018},
  publisher={Taylor \& Francis}
}

@article{wei2020zero,
  title={A zero-inflated gamma model for deconvolved calcium imaging traces},
  author={Wei, Xue-Xin and Zhou, Ding and Grosmark, Andres and Ajabi, Zaki and Sparks, Fraser and Zhou, Pengcheng and Brandon, Mark and Losonczy, Attila and Paninski, Liam},
  journal={arXiv preprint arXiv:2006.03737},
  year={2020}
}

@article{mishchenko2011efficient,
  title={Efficient methods for sampling spike trains in networks of coupled neurons},
  author={Mishchenko, Yuriy and Paninski, Liam},
  journal={The Annals of Applied Statistics},
  pages={1893--1919},
  year={2011},
  publisher={JSTOR}
}


@article{wei2019sequential,
  title={Sequential bayesian detection of spike activities from fluorescence observations},
  author={Wei, Zhuangkun and Li, Bin and Guo, Weisi and Hu, Wenxiu and Zhao, Chenglin},
  journal={IEEE Transactions on Molecular, Biological and Multi-Scale Communications},
  volume={5},
  number={1},
  pages={3--18},
  year={2019},
  publisher={IEEE}
}

@article{sebastian2019spike,
  title={Spike Estimation From Fluorescence Signals Using High-Resolution Property of Group Delay},
  author={Sebastian, Jilt and Kumar, Mari Ganesh and Viraraghavan, Venkata Subramanian and Sur, Mriganka and Murthy, Hema A},
  journal={IEEE Transactions on Signal Processing},
  volume={67},
  number={11},
  pages={2923--2936},
  year={2019},
  publisher={IEEE}
}

@article{ahrens2013whole,
  title={Whole-brain functional imaging at cellular resolution using light-sheet microscopy},
  author={Ahrens, Misha B and Orger, Michael B and Robson, Drew N and Li, Jennifer M and Keller, Philipp J},
  journal={Nature methods},
  volume={10},
  number={5},
  pages={413--420},
  year={2013},
  publisher={Nature Publishing Group}
}


@article{shibue2020deconvolution,
  title={Deconvolution of calcium imaging data using marked point processes},
  author={Shibue, Ryohei and Komaki, Fumiyasu},
  journal={PLoS computational biology},
  volume={16},
  number={3},
  pages={e1007650},
  year={2020},
  publisher={Public Library of Science San Francisco, CA USA}
}

@article{sebastian2021signal,
  title={Signal-to-signal neural networks for improved spike estimation from calcium imaging data},
  author={Sebastian, Jilt and Sur, Mriganka and Murthy, Hema A and Magimai-Doss, Mathew},
  journal={PLoS Computational Biology},
  volume={17},
  number={3},
  pages={e1007921},
  year={2021},
  publisher={Public Library of Science San Francisco, CA USA}
}

@article{im2019importance,
  title={Importance Weighted Adversarial Variational Autoencoders for Spike Inference from calcium Imaging Data},
  author={Im, Daniel Jiwoong and Prakhya, Sridhama and Yan, Jinyao and Turaga, Srinivas and Branson, Kristin},
  journal={arXiv preprint arXiv:1906.03214},
  year={2019}
}

@article{stern2020inferring,
  title={Inferring the spiking rate of a population of neurons from wide-field calcium imaging},
  author={Stern, Merav and Shea-Brown, Eric and Witten, Daniela},
  journal={bioRxiv},
  year={2020},
  publisher={Cold Spring Harbor Laboratory}
}

@article{tsunoda2010statistical,
  title={Statistical Calibration Method for Physiological Ca2+ Fluorescence Signals},
  author={Tsunoda, Takamasa and Oda, Yoshiaki and Omori, Toshiaki and Okada, Masato and Inoue, Masashi and Miyakawa, Hiroyoshi and Aonishi, Toru},
  journal={Australian Journal of Intelligent Information Processing Systems},
  volume={11},
  number={1},
  year={2010}
}
@article{kottas2012bayesian,
  title={Bayesian nonparametric analysis of neuronal intensity rates},
  author={Kottas, Athanasios and Behseta, Sam and Moorman, David E and Poynor, Valerie and Olson, Carl R},
  journal={Journal of Neuroscience Methods},
  volume={203},
  number={1},
  pages={241--253},
  year={2012},
  publisher={Elsevier}
}

@article{ranganathan2010optical,
  title={Optical recording of neuronal spiking activity from unbiased populations of neurons with high spike detection efficiency and high temporal precision},
  author={Ranganathan, Gayathri N and Koester, Helmut J},
  journal={Journal of neurophysiology},
  volume={104},
  number={3},
  pages={1812--1824},
  year={2010},
  publisher={American Physiological Society Bethesda, MD}
}

@article{lindsten2014particle,
  title={Particle Gibbs with ancestor sampling},
  author={Lindsten, Fredrik and Jordan, Michael I and Schon, Thomas B},
  journal={Journal of Machine Learning Research},
  volume={15},
  pages={2145--2184},
  year={2014},
  publisher={MICROTOME PUBL}
}

@article{andrieu2010particle,
  title={Particle markov chain monte carlo methods},
  author={Andrieu, Christophe and Doucet, Arnaud and Holenstein, Roman},
  journal={Journal of the Royal Statistical Society: Series B (Statistical Methodology)},
  volume={72},
  number={3},
  pages={269--342},
  year={2010},
  publisher={Wiley Online Library}
}

@article{park2013quantifying,
  title={Quantifying bursting neuron activity from calcium signals using blind deconvolution},
  author={Park, In Jun and Bobkov, Yuriy V and Ache, Barry W and Principe, Jose C},
  journal={Journal of neuroscience methods},
  volume={218},
  number={2},
  pages={196--205},
  year={2013},
  publisher={Elsevier}
}

@article{pnevmatikakis2016simultaneous,
  title={Simultaneous denoising, deconvolution, and demixing of calcium imaging data},
  author={Pnevmatikakis, Eftychios A and Soudry, Daniel and Gao, Yuanjun and Machado, Timothy A and Merel, Josh and Pfau, David and Reardon, Thomas and Mu, Yu and Lacefield, Clay and Yang, Weijian and others},
  journal={Neuron},
  volume={89},
  number={2},
  pages={285--299},
  year={2016},
  publisher={Elsevier}
}

@inproceedings{dyer2013robust,
  title={A robust and efficient method to recover neural events from noisy and corrupted data},
  author={Dyer, Eva L and Studer, Christoph and Robinson, Jacob T and Baraniuk, Richard G},
  booktitle={2013 6th International IEEE/EMBS Conference on Neural Engineering (NER)},
  pages={593--596},
  year={2013},
  organization={IEEE}
}

@article{kazemipour2017fast,
  title={Fast and stable signal deconvolution via compressible state-space models},
  author={Kazemipour, Abbas and Liu, Ji and Solarana, Krystyna and Nagode, Daniel A and Kanold, Patrick O and Wu, Min and Babadi, Behtash},
  journal={IEEE Transactions on Biomedical Engineering},
  volume={65},
  number={1},
  pages={74--86},
  year={2017},
  publisher={IEEE}
}

@inproceedings{dyer2010recovering,
  title={Recovering spikes from noisy neuronal calcium signals via structured sparse approximation},
  author={Dyer, Eva L and Duarte, Marco F and Johnson, Don H and Baraniuk, Richard G},
  booktitle={International Conference on Latent Variable Analysis and Signal Separation},
  pages={604--611},
  year={2010},
  organization={Springer}
}

@article{quan2010method,
  title={Method to reconstruct neuronal action potential train from two-photon calcium imaging},
  author={Quan, Tingwei and Liu, Xiuli and Lv, Xiaohua and Chen, Wei R and Zeng, Shaoqun},
  journal={Journal of biomedical optics},
  volume={15},
  number={6},
  pages={066002},
  year={2010},
  publisher={International Society for Optics and Photonics}
}

@article{theis2013beyond,
  title={Beyond GLMs: a generative mixture modeling approach to neural system identification},
  author={Theis, Lucas and Chagas, Andr{\`e} Maia and Arnstein, Daniel and Schwarz, Cornelius and Bethge, Matthias},
  journal={PLoS computational biology},
  volume={9},
  number={11},
  pages={e1003356},
  year={2013},
  publisher={Public Library of Science San Francisco, USA}
}

@article{tsunoda2010estimation,
  title={Estimation of intracellular calcium ion concentration by nonlinear state space modeling and expectation-maximization algorithm for parameter estimation},
  author={Tsunoda, Takamasa and Omori, Toshiaki and Miyakawa, Hiroyoshi and Okada, Masato and Aonishi, Toru},
  journal={Journal of the Physical Society of Japan},
  volume={79},
  number={12},
  pages={124801},
  year={2010},
  publisher={The Physical Society of Japan}
}
@article{vogelstein2009spike,
  title={Spike inference from calcium imaging using sequential Monte Carlo methods},
  author={Vogelstein, Joshua T and Watson, Brendon O and Packer, Adam M and Yuste, Rafael and Jedynak, Bruno and Paninski, Liam},
  journal={Biophysical journal},
  volume={97},
  number={2},
  pages={636--655},
  year={2009},
  publisher={Elsevier}
}

@article{pnevmatikakis2014structured,
  title={A structured matrix factorization framework for large scale calcium imaging data analysis},
  author={Pnevmatikakis, Eftychios A and Gao, Yuanjun and Soudry, Daniel and Pfau, David and Lacefield, Clay and Poskanzer, Kira and Bruno, Randy and Yuste, Rafael and Paninski, Liam},
  journal={arXiv preprint arXiv:1409.2903},
  year={2014}
}

@article{poskanzer2014structured,
  title={A structured matrix factorization framework for large scale calcium imaging data analysis},
  author={Poskanzer, Kira and Pnevmatikakis, EA and Gao, Y and Soudry, D and Pfau, D and Lacefield, C and Bruno, R and Yuste, R and Paninski, L},
  year={2014}
}

@article{speiser2017fast,
  title={Fast amortized inference of neural activity from calcium imaging data with variational autoencoders},
  author={Speiser, Artur and Yan, Jinyao and Archer, Evan and Buesing, Lars and Turaga, Srinivas C and Macke, Jakob H},
  journal={arXiv preprint arXiv:1711.01846},
  year={2017}
}

@article{sasaki2008fast,
  title={Fast and accurate detection of action potentials from somatic calcium fluctuations},
  author={Sasaki, Takuya and Takahashi, Naoya and Matsuki, Norio and Ikegaya, Yuji},
  journal={Journal of neurophysiology},
  volume={100},
  number={3},
  pages={1668--1676},
  year={2008},
  publisher={American Physiological Society}
}

@article{rahmati2016inferring,
  title={Inferring neuronal dynamics from calcium imaging data using biophysical models and Bayesian inference},
  author={Rahmati, Vahid and Kirmse, Knut and Markovi{\'c}, Dimitrije and Holthoff, Knut and Kiebel, Stefan J},
  journal={PLoS computational biology},
  volume={12},
  number={2},
  pages={e1004736},
  year={2016},
  publisher={Public Library of Science San Francisco, CA USA}
}

@article{friedrich2016fast,
  title={Fast active set methods for online spike inference from calcium imaging},
  author={Friedrich, Johannes and Paninski, Liam},
  journal={Advances In Neural Information Processing Systems},
  volume={29},
  pages={1984--1992},
  year={2016}
}

@article{malik2011denoising,
  title={Denoising two-photon calcium imaging data},
  author={Malik, Wasim Q and Schummers, James and Sur, Mriganka and Brown, Emery N},
  journal={PloS one},
  volume={6},
  number={6},
  pages={e20490},
  year={2011},
  publisher={Public Library of Science San Francisco, USA}
}

@article{fletcher2014scalable,
  title={Scalable inference for neuronal connectivity from calcium imaging},
  author={Fletcher, Alyson K and Rangan, Sundeep},
  journal={arXiv preprint arXiv:1409.0289},
  year={2014}
}

@article{jewell2020fast,
  title={Fast nonconvex deconvolution of calcium imaging data},
  author={Jewell, Sean W and Hocking, Toby Dylan and Fearnhead, Paul and Witten, Daniela M},
  journal={Biostatistics},
  volume={21},
  number={4},
  pages={709--726},
  year={2020},
  publisher={Oxford University Press}
}
@article{mishchencko2011bayesian,
  title={A Bayesian approach for inferring neuronal connectivity from calcium fluorescent imaging data},
  author={Mishchencko, Yuriy and Vogelstein, Joshua T and Paninski, Liam},
  journal={The Annals of Applied Statistics},
  pages={1229--1261},
  year={2011},
  publisher={JSTOR}
}

@article{huys2009smoothing,
  title={Smoothing of, and parameter estimation from, noisy biophysical recordings},
  author={Huys, Quentin JM and Paninski, Liam},
  journal={PLoS computational biology},
  volume={5},
  number={5},
  pages={e1000379},
  year={2009},
  publisher={Public Library of Science San Francisco, USA}
}

@article{onativia2013finite,
  title={A finite rate of innovation algorithm for fast and accurate spike detection from two-photon calcium imaging},
  author={O{\~n}ativia, Jon and Schultz, Simon R and Dragotti, Pier Luigi},
  journal={Journal of neural engineering},
  volume={10},
  number={4},
  pages={046017},
  year={2013},
  publisher={IOP Publishing}
}

@article{onativia2015sparse,
  title={Sparse sampling: theory, methods and an application in neuroscience},
  author={O{\~n}ativia, Jon and Dragotti, Pier Luigi},
  journal={Biological cybernetics},
  volume={109},
  number={1},
  pages={125--139},
  year={2015},
  publisher={Springer}
}

@article{zhou2018efficient,
  title={Efficient and accurate extraction of in vivo calcium signals from microendoscopic video data},
  author={Zhou, Pengcheng and Resendez, Shanna L and Rodriguez-Romaguera, Jose and Jimenez, Jessica C and Neufeld, Shay Q and Giovannucci, Andrea and Friedrich, Johannes and Pnevmatikakis, Eftychios A and Stuber, Garret D and Hen, Rene and others},
  journal={Elife},
  volume={7},
  pages={e28728},
  year={2018},
  publisher={eLife Sciences Publications Limited}
}


@article{friedrich2017fast,
  title={Fast online deconvolution of calcium imaging data},
  author={Friedrich, Johannes and Zhou, Pengcheng and Paninski, Liam},
  journal={PLoS computational biology},
  volume={13},
  number={3},
  pages={e1005423},
  year={2017},
  publisher={Public Library of Science San Francisco, CA USA}
}

@inproceedings{pnevmatikakis2013bayesian,
  title={Bayesian spike inference from calcium imaging data},
  author={Pnevmatikakis, Eftychios A and Merel, Josh and Pakman, Ari and Paninski, Liam},
  booktitle={2013 Asilomar Conference on Signals, Systems and Computers},
  pages={349--353},
  year={2013},
  organization={IEEE}
}

@article{jewell2018exact,
  title={Exact spike train inference via $\ell_0$ optimization},
  author={Jewell, Sean and Witten, Daniela},
  journal={The annals of applied statistics},
  volume={12},
  number={4},
  pages={2457},
  year={2018},
  publisher={NIH Public Access}
}

@article{kerr2005imaging,
  title={Imaging input and output of neocortical networks in vivo},
  author={Kerr, Jason ND and Greenberg, David and Helmchen, Fritjof},
  journal={Proceedings of the National Academy of Sciences},
  volume={102},
  number={39},
  pages={14063--14068},
  year={2005},
  publisher={National Acad Sciences}
}

@article{yaksi2006reconstruction,
  title={Reconstruction of firing rate changes across neuronal populations by temporally deconvolved Ca 2+ imaging},
  author={Yaksi, Emre and Friedrich, Rainer W},
  journal={Nature methods},
  volume={3},
  number={5},
  pages={377--383},
  year={2006},
  publisher={Nature Publishing Group}
}

@article{holekamp2008fast,
  title={Fast three-dimensional fluorescence imaging of activity in neural populations by objective-coupled planar illumination microscopy},
  author={Holekamp, Terrence F and Turaga, Diwakar and Holy, Timothy E},
  journal={Neuron},
  volume={57},
  number={5},
  pages={661--672},
  year={2008},
  publisher={Elsevier}
}
@article{berens2018community,
  title={Community-based benchmarking improves spike rate inference from two-photon calcium imaging data},
  author={Berens, Philipp and Freeman, Jeremy and Deneux, Thomas and Chenkov, Nikolay and McColgan, Thomas and Speiser, Artur and Macke, Jakob H and Turaga, Srinivas C and Mineault, Patrick and Rupprecht, Peter and others},
  journal={PLoS computational biology},
  volume={14},
  number={5},
  pages={e1006157},
  year={2018},
  publisher={Public Library of Science San Francisco, CA USA}
}

@article{d2021bayesian,
  title={Bayesian nonparametric analysis for the detection of spikes in noisy calcium imaging data},
  author={D'Angelo, Laura and Canale, Antonio and Yu, Zhaoxia and Guindani, Michele},
  journal={arXiv preprint arXiv:2102.09403},
  year={2021}
}

@article{rupprecht2021database,
  title={A database and deep learning toolbox for noise-optimized, generalized spike inference from calcium imaging},
  author={Rupprecht, Peter and Carta, Stefano and Hoffmann, Adrian and Echizen, Mayumi and Blot, Antonin and Kwan, Alex C and Dan, Yang and Hofer, Sonja B and Kitamura, Kazuo and Helmchen, Fritjof and others},
  journal={Nature Neuroscience},
  volume={24},
  number={9},
  pages={1324--1337},
  year={2021},
  publisher={Nature Publishing Group}
}

@article{hoang2020improved,
  title={Improved hyperacuity estimation of spike timing from calcium imaging},
  author={Hoang, Huu and Sato, Masa-aki and Shinomoto, Shigeru and Tsutsumi, Shinichiro and Hashizume, Miki and Ishikawa, Tomoe and Kano, Masanobu and Ikegaya, Yuji and Kitamura, Kazuo and Kawato, Mitsuo and others},
  journal={Scientific reports},
  volume={10},
  number={1},
  pages={1--16},
  year={2020},
  publisher={Nature Publishing Group}
}

@article{mukamel2009automated,
  title={Automated analysis of cellular signals from large-scale calcium imaging data},
  author={Mukamel, Eran A and Nimmerjahn, Axel and Schnitzer, Mark J},
  journal={Neuron},
  volume={63},
  number={6},
  pages={747--760},
  year={2009},
  publisher={Elsevier}
}

@article{grewe2010high,
  title={High-speed in vivo calcium imaging reveals neuronal network activity with near-millisecond precision},
  author={Grewe, Benjamin F and Langer, Dominik and Kasper, Hansj{\"o}rg and Kampa, Bj{\"o}rn M and Helmchen, Fritjof},
  journal={Nature methods},
  volume={7},
  number={5},
  pages={399--405},
  year={2010},
  publisher={Nature Publishing Group}
}

@article{vogelstein2010fast,
  title={Fast nonnegative deconvolution for spike train inference from population calcium imaging},
  author={Vogelstein, Joshua T and Packer, Adam M and Machado, Timothy A and Sippy, Tanya and Babadi, Baktash and Yuste, Rafael and Paninski, Liam},
  journal={Journal of neurophysiology},
  volume={104},
  number={6},
  pages={3691--3704},
  year={2010},
  publisher={American Physiological Society Bethesda, MD}
}

@article{stetter2012model,
  title={Model-free reconstruction of excitatory neuronal connectivity from calcium imaging signals},
  author={Stetter, Olav and Battaglia, Demian and Soriano, Jordi and Geisel, Theo},
  year={2012},
  publisher={Public Library of Science San Francisco, USA}
}

@article{zhang2023fast,
  title={Fast and sensitive GCaMP calcium indicators for imaging neural populations},
  author={Zhang, Yan and R{\'o}zsa, M{\'a}rton and Liang, Yajie and Bushey, Daniel and Wei, Ziqiang and Zheng, Jihong and Reep, Daniel and Broussard, Gerard Joey and Tsang, Arthur and Tsegaye, Getahun and others},
  journal={Nature},
  volume={615},
  number={7954},
  pages={884--891},
  year={2023},
  publisher={Nature Publishing Group UK London}
}

@article{theis2016benchmarking,
  title={Benchmarking spike rate inference in population calcium imaging},
  author={Theis, Lucas and Berens, Philipp and Froudarakis, Emmanouil and Reimer, Jacob and Ros{\'o}n, Miroslav Rom{\'a}n and Baden, Tom and Euler, Thomas and Tolias, Andreas S and Bethge, Matthias},
  journal={Neuron},
  volume={90},
  number={3},
  pages={471--482},
  year={2016},
  publisher={Elsevier}
}

@article{pachitariu2017suite2p,
  title={Suite2p: beyond 10,000 neurons with standard two-photon microscopy},
  author={Pachitariu, Marius and Stringer, Carsen and Dipoppa, Mario and Schr{\"o}der, Sylvia and Rossi, L Federico and Dalgleish, Henry and Carandini, Matteo and Harris, Kenneth D},
  journal={BioRxiv},
  year={2017},
  publisher={Biorxiv}
}


@article{giovannucci2019caiman,
  title={CaImAn an open source tool for scalable calcium imaging data analysis},
  author={Giovannucci, Andrea and Friedrich, Johannes and Gunn, Pat and Kalfon, J{\'e}r{\'e}mie and Brown, Brandon L and Koay, Sue Ann and Taxidis, Jiannis and Najafi, Farzaneh and Gauthier, Jeffrey L and Zhou, Pengcheng and others},
  journal={Elife},
  volume={8},
  pages={e38173},
  year={2019},
  publisher={eLife Sciences Publications Limited}
}

@article{lutcke2013inference,
  title={Inference of neuronal network spike dynamics and topology from calcium imaging data},
  author={L{\"u}tcke, Henry and Gerhard, Felipe and Zenke, Friedemann and Gerstner, Wulfram and Helmchen, Fritjof},
  journal={Frontiers in neural circuits},
  volume={7},
  pages={201},
  year={2013},
  publisher={Frontiers}
}

@article{deneux2016accurate,
  title={Accurate spike estimation from noisy calcium signals for ultrafast three-dimensional imaging of large neuronal populations in vivo},
  author={Deneux, Thomas and Kaszas, Attila and Szalay, Gergely and Katona, Gergely and Lakner, Tam{\'a}s and Grinvald, Amiram and R{\'o}zsa, Bal{\'a}zs and Vanzetta, Ivo},
  journal={Nature communications},
  volume={7},
  number={1},
  pages={1--17},
  year={2016},
  publisher={Nature Publishing Group}
}
@article{chen2013overview,
  title={An overview of bayesian methods for neural spike train analysis},
  author={Chen, Zhe},
  journal={Computational intelligence and neuroscience},
  volume={2013},
  year={2013},
  publisher={Hindawi}
}

@article{greenberg2018accurate,
  title={Accurate action potential inference from a calcium sensor protein through biophysical modeling},
  author={Greenberg, David S and Wallace, Damian J and Voit, Kay-Michael and Wuertenberger, Silvia and Czubayko, Uwe and Monsees, Arne and Handa, Takashi and Vogelstein, Joshua T and Seifert, Reinhard and Groemping, Yvonne and others},
  journal={BioRxiv},
  pages={479055},
  year={2018},
  publisher={Cold Spring Harbor Laboratory}
}
